# Linear Mixed-effects model to quantify the association between somatic cell count and milk production in Italian dairy herds

**DOI:** 10.1101/2022.07.15.500192

**Authors:** Tiantian Luo, Wilma Steeneveld, Mirjam Nielen, Lucio Zanini, Alfonso Zecconi

## Abstract

Milk production loss due to mastitis in dairy herds is economically important. Before estimating the economic impacts of mastitis, it is crucial to quantify the association between mastitis and milk production. The objective of this study was to estimate the association between somatic cell count (SCC, as an indicator of intramammary infection due to mastitis) and milk production for dairy cows in Lombardy (Italy). The test-day (TD) records data of 3,816 dairy herds located in 3 different geographical areas of Lombardy from January 2016 to December 2018 were used. After data editing, the final dataset comprised 10,445,464 TD records from 2,970 farms and 826,831 cows. The analysis was carried out by using a mixed-effects model with seven fixed effects (geographical area, breed, days in milk, parity, season and year) and nested random effects for each cow and herd. The results indicated that the SCC had a negative association with milk production. When the SCC increased every two-fold, the milk production lost 0.830 (95% CI: −0.832, −0.828) kg/cow/day in the whole Lombardy. These results can be used for economic calculations on the costs of mastitis

## Introduction

Bovine mastitis is one of the most frequently occurring and costly diseases that affects the welfare of dairy cows [1–3]. The total costs of mastitis include production losses, prevention and treatment costs, culling, changes in product quality and the risk of other diseases [4]. It is known that these costs can vary a lot between farms [5–7]. Van Soest et al. (2016) reported that the average total cost of mastitis can achieve €240/lactating cow per year. In Italy, the average cost of a single clinical mastitis case was estimated at €177[8] and a decrease of average milk yield per cow of around 2 kg/d was estimated in herds with contagious pathogens when compared with contagious-free herds [9].

To estimate the economic impacts of mastitis, it is crucial to quantify the production losses for both clinical and subclinical mastitis (SCM), since production losses are considered as one of the largest components of total costs of mastitis [7]. The production losses due to SCM can be determined with the Somatic Cell Count (SCC) [10]. SCC is one of the standard diagnostic tests to detect SCM, and at most farms monthly measured at cow level as part of the milk recording system. A value higher than 200,000 cells/ml is considered positive for SCM[11,12]. An increased SCC is thus an indication of an inflammatory reaction, which can substantially reduce milk production. Milk production loss associated with increasing SCC was investigated by several studies[13–16]. Halasa et al. (2009) reported that when the SCC increased 2-fold after a low SCC period, the milk production decreased by 0.38 and 0.46 kg/d for primiparous and multiparous cows, respectively. Hand et al. (2012) found that the daily milk loss ranged from 0.35 to 4.70kg for SCC values from 200,000 cells/ml to 2,000,000 cells/ml, and the whole lactation milk loss ranged from 165 to 919kg per lactation. In Brazil, Gonçalves et al. (2018) revealed that milk losses per unit increase of log-transformed SCC varied between 0.55 kg/d and 2.45 kg/d in 3 different lactation stages and 3 parities.

The association between SCC and milk production was thus estimated, but mostly on relatively large scale intensive dairy herds, and specific management in North-West European and American circumstances [16–18]. This association is much less determined for small scale dairy herds or in areas where dairy herds are less intensive such as in mountain areas. Furthermore, it has not been quantified in the Lombardy region of Italy. Lombardy represents the leading milk-producing region in Italy with more than 40% of the national production, and the dairy herds in the region here are in different geographical areas (the Alps, Sub-Alps and Po valley) with both large scales intensive dairy herds and small-scale dairy herds. The objective of this study was to estimate the association between SCC and milk production for dairy cows in Lombardy, due to the importance of milk production in this area and the absence of specific recent studies on the association. The results could help in prioritizing the interventions[9] from the advisory services (i.e. from regional breeder association). They may be also used as a reference for areas in other countries with similar characteristics as Lombardy (i.e. other European countries, south America).

## Materials and Method

### Milk sampling and analysis

The data to be analyzed includes all the herds in Lombardy associated with Italian Breeder Association (AIA) and applying routine milk record sampling for three years. Individual cow samplings were performed by certified methods currently applied by the AIA at the laboratories of Regional Breeders Association of Lombardy (ARAL). Samples were taken about every 5 weeks, delivered refrigerated to ARAL labs the same day, and analyzed within 30 h from sampling. SCC was performed by certified methods, currently applied by AIA at the laboratories of ARAL on Fossomatic FC (Foss DK). Cow and Milk Test Records (MTR) were supplied by AIA through ARAL and they were: herdID, cowID, number of lactations, SCC and milk yield at every milk test conducted.

These data were collected from nine provinces including all the three main geographical areas of the region: Alps, sub-Alps and Po valley. The initial data included 3,816 herds and 905,119 cows with 11,396,685 test-day (TD) records from January 2016 to December 2018.

### Data editing

The procedure of data editing is presented in Fig 1. First, TD records with missing values on SCC were excluded. Secondly, the herds with a size less than 30 cows were excluded in sub-Alps and Po valley. In the Alps area, the herds with less than 10 cows were excluded. Thirdly, the TD records with SCC higher than 9,999,000 cells/mL were excluded. Fourthly, TD records with days in milk (DIM) longer than 400 days were excluded. Finally, if the cow only had one test in a lactation, these records were also excluded. The final dataset comprised 10,445,464 TD records from 2,970 farms and 826,831 cows.

**Fig 1.**
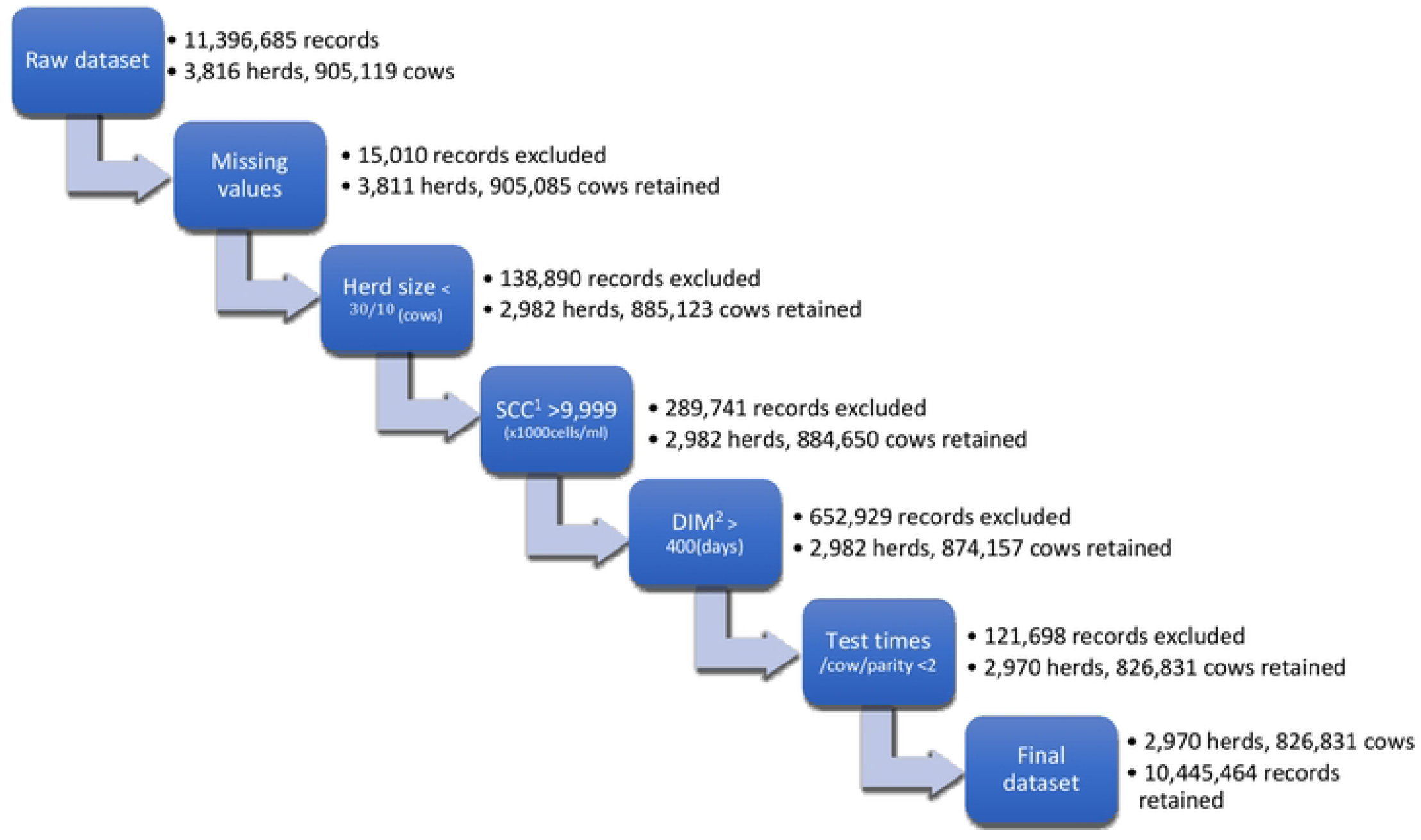
Editing criteria and number of herds, cows and test day records retained and excluded ^*1*^*Somatic Cell Count*. ^*2*^*Days in Milk*.

The geographical areas of the herd location were categorized into 3 groups: Alps, sub-Alps and Po valley. There were 24 types of breeds, including Holstein, Brown Swiss, Simmrnyhsl, Jersey, local breeds and mixed breeds. In Po valley, 94.2% of TD records were from Holstein cows. 1.0% and 4.8% of records were from Brown Swiss and other breeds, respectively. Similarly in Sub-Alps, 85.8%, 7.1% and 7.2% of TD records were from Holstein, Brown Swiss and other breeds, respectively. In the Alps, Brown Swiss was the dominant breed which was 42.8% of the whole TD records. 35.2% and 22.0% of TD records were from Holstein and other breeds, respectively. Because some of the breeds have a higher capability of milk production and tend to have higher SCC [10], the breeds were categorized into 3 groups: Holstein, Brown Swiss and other breeds. The DIM were categorized into 13 stages with the interval of every 30 days. The parity was categorized into 3 groups: parity 1, parity 2 and parity equals or greater than 3. According to the date of each TD record, 4 season groups (April, May and June as spring, July, August and September as summer, October, November and December as autumn, January, February and March as winter) and 3 year-groups (2016, 2017 and 2018) were categorized.

### Statistical analysis

Given the multilevel structure of the longitudinal data, a linear mixed-effects model was used to estimate the association between SCC and milk production. SCC is a right-skewed variable, and this violated the assumption of the linear model. Therefore, Log_2_(SCC) (usually Log_2_(SCC) defined as Somatic cell score, SCS) was used to fit in the linear model [19]. Several explanatory variables might also influence milk production, and were inserted in the model as fixed effects. These fixed effects included Area (categorical, 3 levels), Breed (categorical, 3 levels), DIM (categorical, 13 levels), Parity (categorical, 3 levels), Season (categorical, 4 levels) and Year (categorical, 3 levels). The TD records were collected during 3 years, which were multiple observations per cow and farm. For an individual cow, the daily milk production values were correlated. Furthermore, the cows within the same herd were correlated. Therefore, the nested random effects (random intercepts) for each cow and herd were introduced into the model. The model applied is as following:

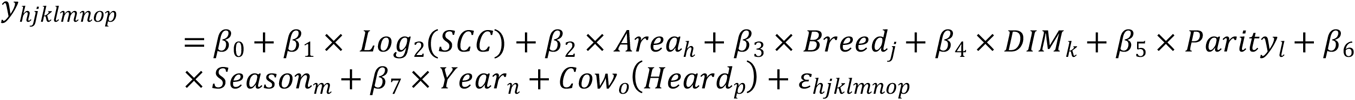

Where *y*_*hjklmnop*_ was the milk production for each cow *o* in herd *p* in the area *h*, the breed *j*, the DIM class *k*, the number of parity *l*, the season *m* and the year *n*; β_0_ was the overall mean of milk production; β_1_ was the regression coefficient of the binary logarithm of the SCC × 10^3^ cells/mL; β_2_ was the regression coefficient of the *h*_*th*_ class of area; β_3_ was the regression coefficient of the *j*_*th*_ class of breed; β_4_ was the regression coefficient of the *k*_*th*_ class of DIM; β_5_ was the regression coefficient of the *l*_*th*_ class of parity; β_6_ was the regression coefficient of the *m*_*th*_ class of season; β_7_ was the regression coefficient of the *n*_*th*_ class of year; *Cow*_*o*_ and *Herd*_*p*_ were the random effects; *e*_*hjklmnop*_ was the residual error. Log_2_(SCC), area, breed, DIM, parity, season and year were the fixed effects of milk production.

Model selection was based on the Akaike information criterion (AIC). Moreover, the model assumptions were checked for multicollinearity with Variance Inflation Factors (VIF), normality of residuals, homoscedasticity, homogeneity of variance and influential observations [20]. For the performance of the mixed models, the marginal R-squared measured the proportion of the variance that was explained by the fixed effects. The conditional R-squared, on the contrary, measured the proportion of the total variance that was explained by both fixed and random effects in the full model. Similar to R-squared, the Intraclass Correlation Coefficients (ICC) also provided information on the explained variance, which was explained by the grouping structure in the population (Hox, 2010). These indexes were all reported after fitting the models.

All computations were performed with *R* (R Core Team, 2020). The restricted maximum likelihood (REML) estimates of the parameters in linear mixed-effects models were determined using the *lmer* function in the *lme4* [21] package. The fit of the model was tested by the *performance* (0.7.0) package. *Dplyr* (0.8.5), *lubridate* (1.7.9)and *ggplot2* (3.3.0) were used for data editing and data visualisation.

## Results

### Descriptive analysis

Descriptive statistics of the final dataset is presented in Table 1. Among Lombardian dairy herds, the median herd size was 176 cows, with an average herd size of 228 cows, and a range from 10 to 1,099 cows. The average herd size in the Alps, Sub-Alps and Po valley were 86, 109 and 233 cows, respectively. The average milk production level at the TD records level was 32.8 kg/d and a range from 0.2 to 109.7 kg/d. The average SCC at the TD records level were 308 × 10^3^ cells/mL, ranging from 1 to 9,999 × 10^3^ cells/mL.

**Table 1.** Descriptive statistics of the final data (10,445,464 test-day records from 826,831 cows on 2,970 herds in Lombardy, including 264,195 test-day records from the Alps, 108,287 test-day records from Sub-Alps and 10,072,982 test-day records from Po valley) at the test-day record level.

|  | Overall |  |  |  |  | Alps |  |  |  |  | Sub-Alps |  |  |  |  | Po valley |  |  |  |  |
| --- | --- | --- | --- | --- | --- | --- | --- | --- | --- | --- | --- | --- | --- | --- | --- | --- | --- | --- | --- | --- |
|  | Min. | Median | Mean | Max. | SD | Min. | Median | Mean | Max. | SD | Min. | Median | Mean | Max. | SD | Min. | Median | Mean | Max. | SD |
| **Herd size (nr of cows)** | 10 | 176 | 228 | 1099 | 179 | 10 | 47 | 86 | 369 | 93 | 30 | 89 | 109 | 269 | 67 | 30 | 179 | 233 | 1099 | 179 |
| **Milk production (kg/d)** | 0.2 | 32.2 | 32.8 | 109.7 | 9.6 | 0.6 | 25.7 | 26.3 | 92.6 | 9.1 | 1.2 | 30.9 | 31.5 | 87.6 | 9.2 | 0.2 | 32.4 | 32.9 | 110 | 9.6 |
| **SCC<sup>1</sup> (x 10<sup>3</sup> cells/mL)** | 1 | 80 | 308 | 9,999 | 779 | 1 | 91 | 319 | 9,990 | 737 | 1 | 73 | 274 | 9,938 | 710 | 1 | 80 | 308 | 9,999 | 780 |
| **Fat (%)** | 2.0 | 3.8 | 3.9 | 7.0 | 0.8 | 2.0 | 4.1 | 4.2 | 7.0 | 0.8 | 2.0 | 3.9 | 3.9 | 7.0 | 0.7 | 2.0 | 3.8 | 3.9 | 7.0 | 0.8 |
| **Protein (%)** | 1.5 | 3.4 | 3.4 | 6.0 | 0.4 | 1.6 | 3.6 | 3.6 | 6.0 | 0.4 | 2.0 | 3.1 | 3.4 | 5.8 | 0.4 | 1.5 | 3.4 | 3.4 | 6.0 | 0.4 |
| **DIM<sup>2</sup> (day)** | 5 | 162 | 170 | 399 | 102 | 5 | 158 | 168 | 399 | 104 | 5 | 162 | 170 | 399 | 103 | 5 | 158 | 168 | 399 | 104 |

<sup>1</sup>Somatic Cell Count.
<sup>2</sup>Days in Milk.

Milk production for different breed groups of cows was presented in Fig 2. The average milk production of Holstein cows was 33.1 kg/d which was the highest among all breed groups. Brown Swiss cows produced 25.6 kg/d milk on average, and other breeds of cows produced 29.3 kg/d milk on average.

**Fig 2.**
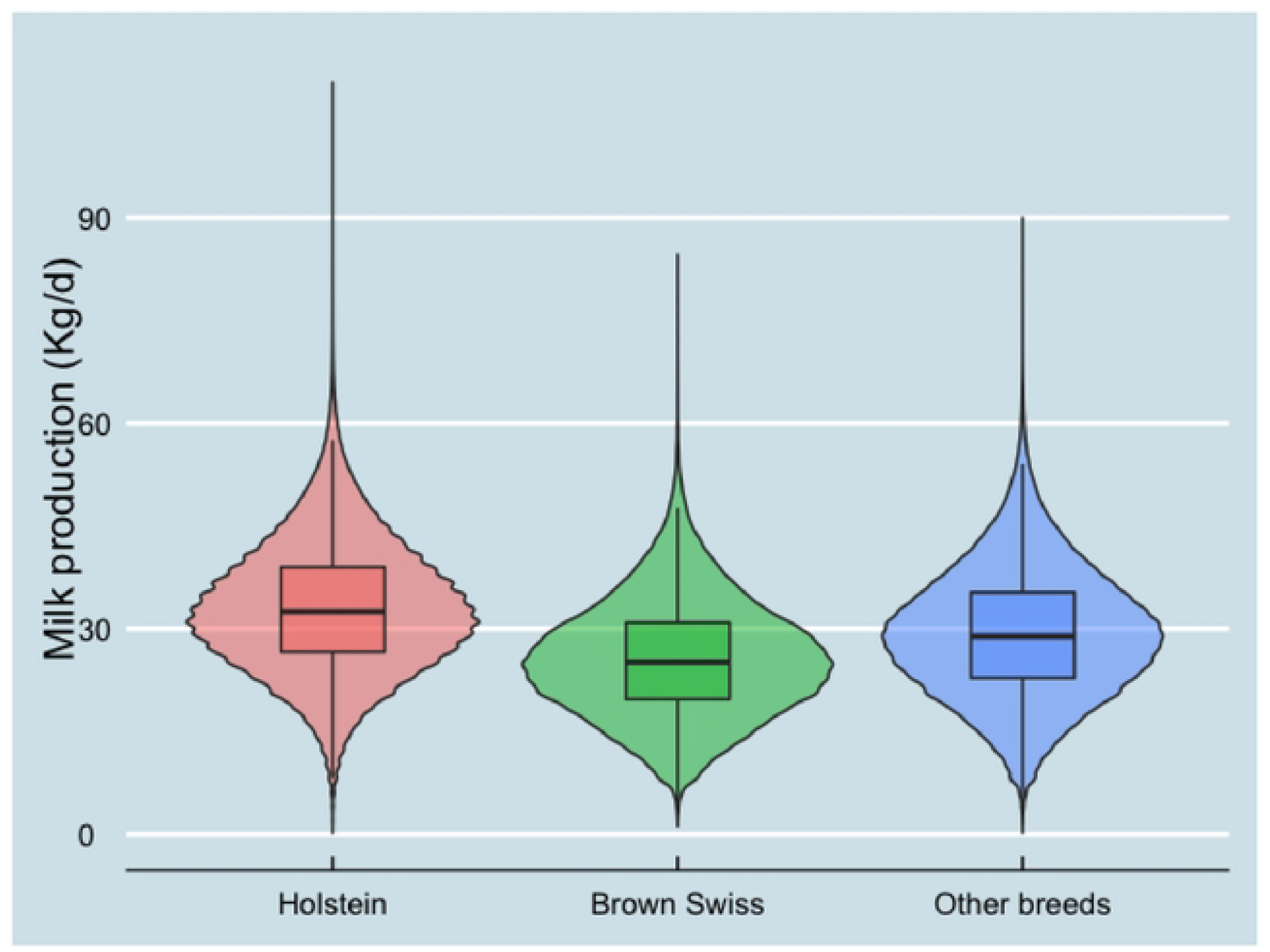
Milk production at Test-day records level in different breed groups (10,445,464 test-day records from 826,831 cows on 2,970 herds)

SCS for different milk production levels and breed groups are presented in Figs 3 and 4. SCS decreased when milk production increased. Holstein cows had the lowest SCS among all breeds.

**Fig 3.**
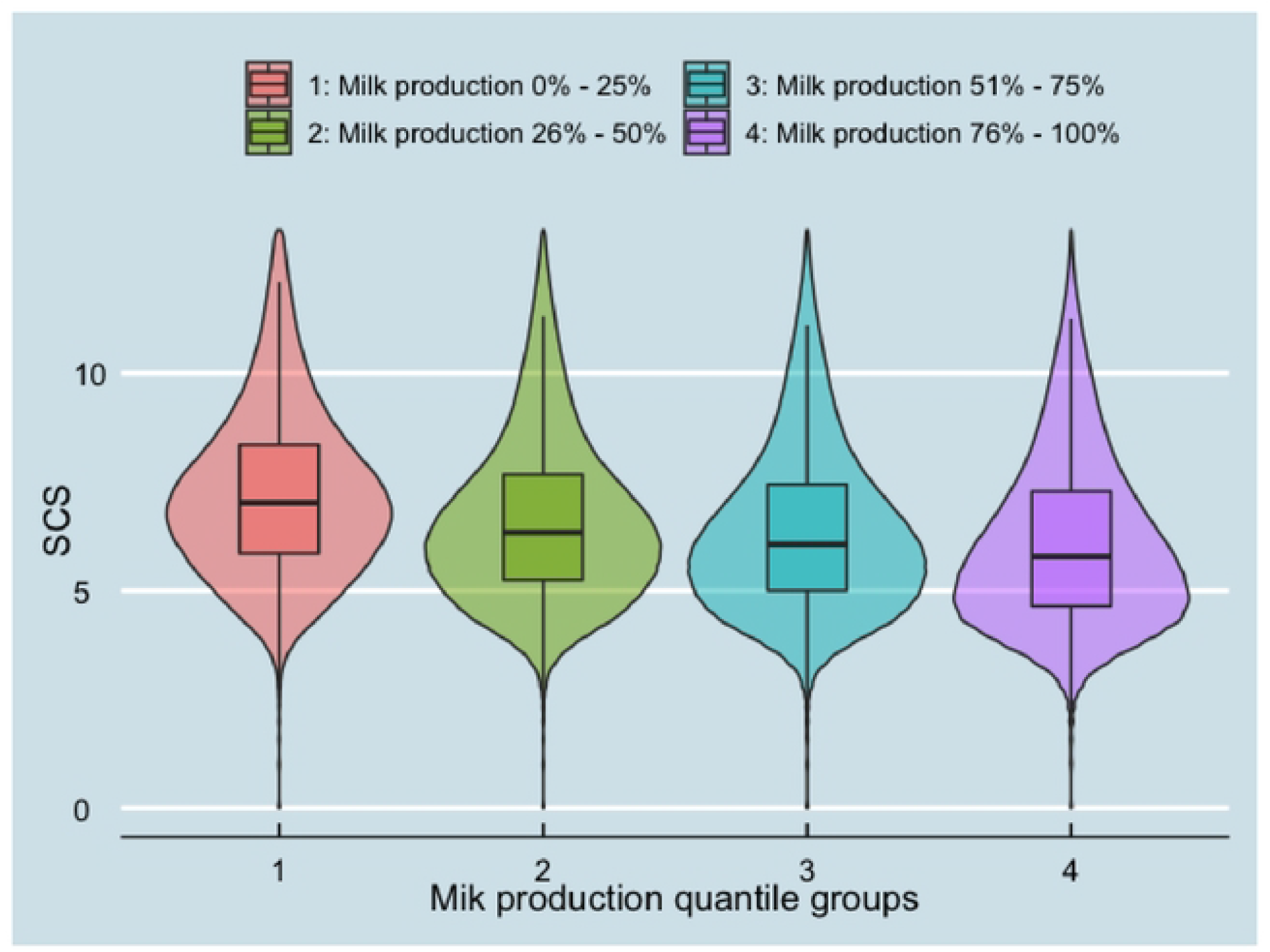
SCS at Test-day records level in different milk production quantile groups (10,445,464 test-day records from 826,831 cows on 2,970 herds)

**Fig 4.**
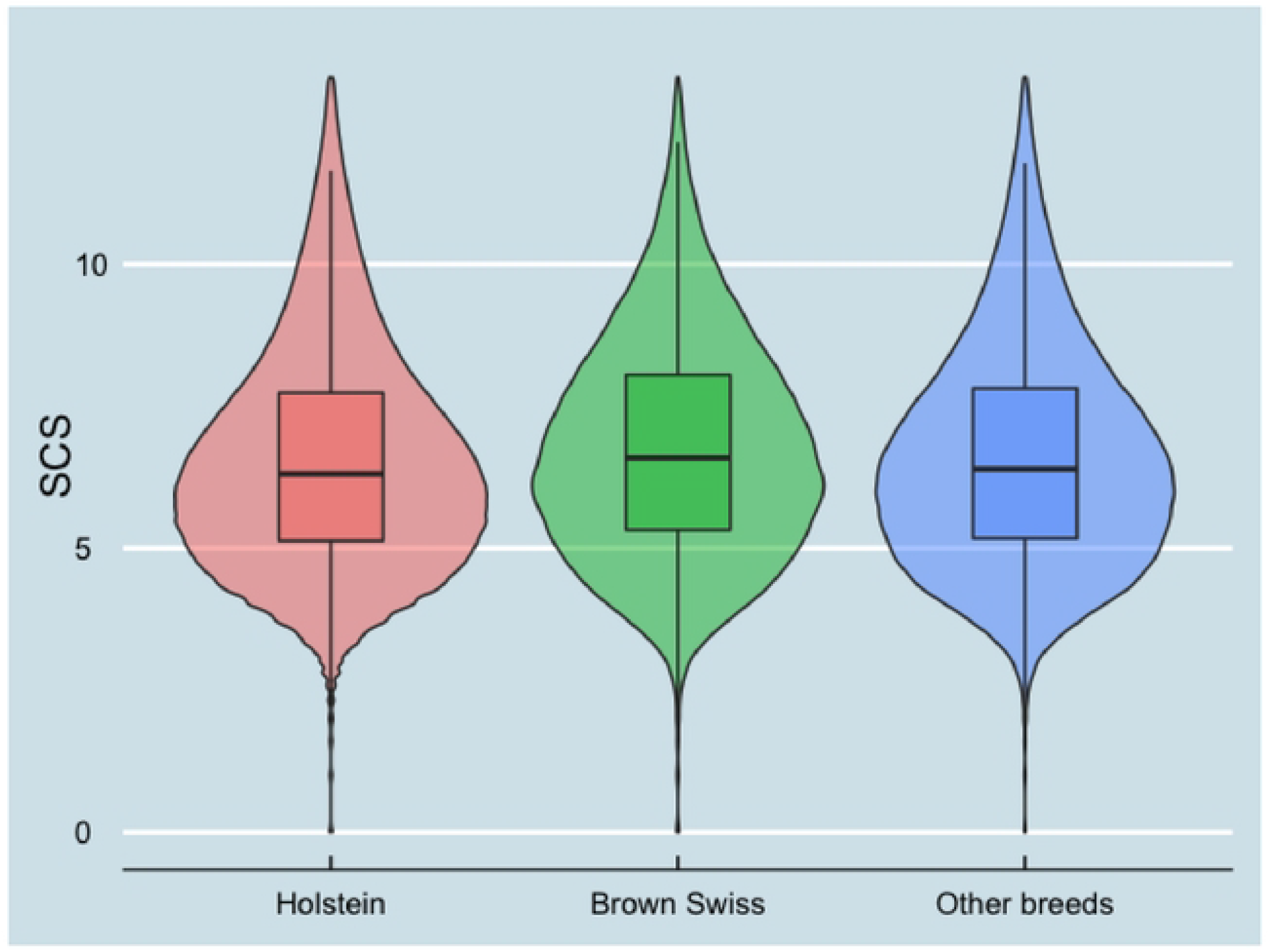
SCS at Test-day records level in different breed groups (10,445,464 test-day records from 826,831 cows on 2,970 herds)

In different areas and cow breeds, the average milk production decreased as SCC increased (see Figs 5 and 6). In the Alps, however, the average milk production decreased more with higher SCC than in other areas. When SCC ≥ 400,000 cells/ml, this decreasing trend of milk production was not obvious amymore.

**Fig 5.**
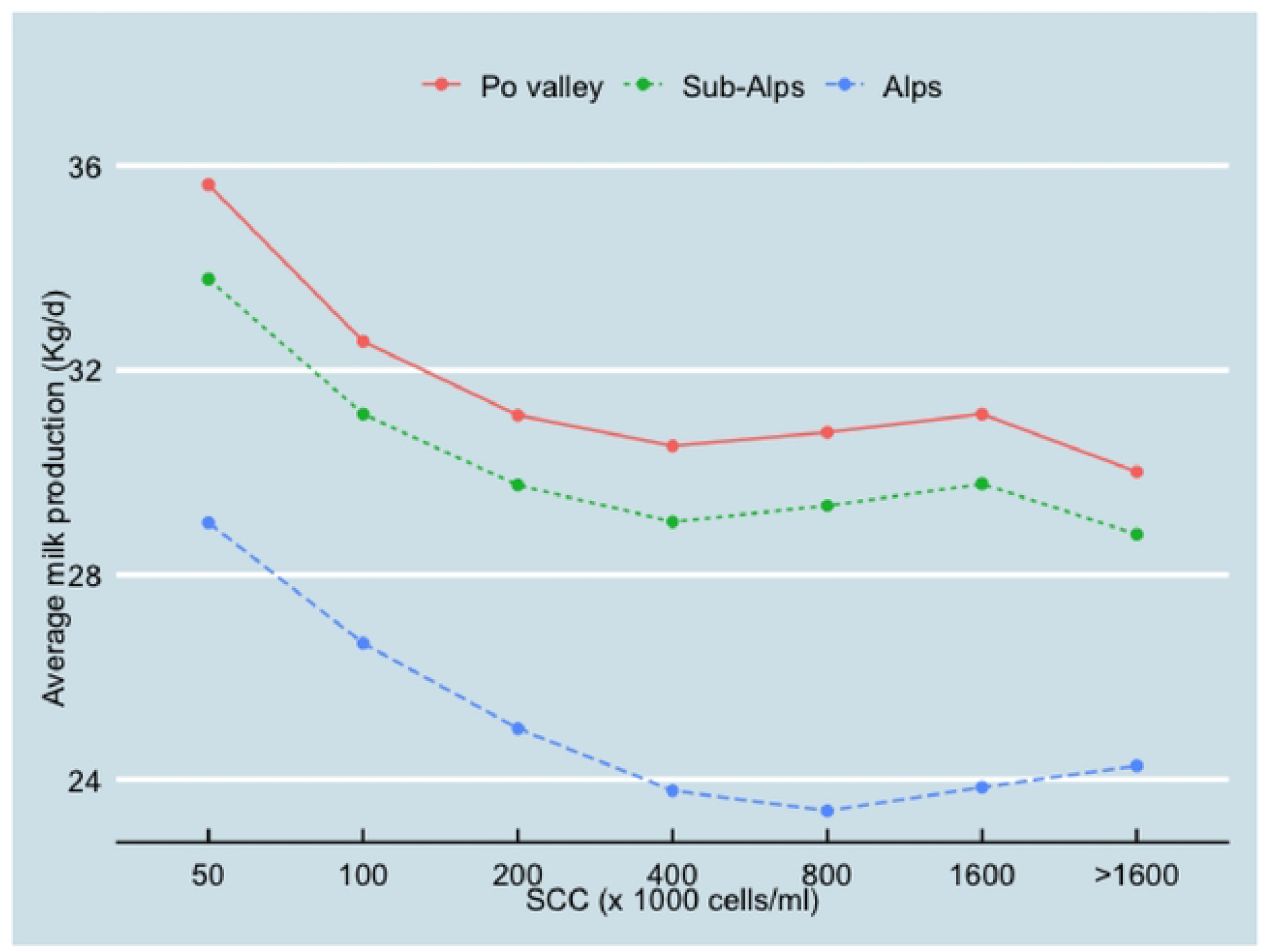
Relationship between SCC and average milk production (10,445,464 test-day records from 826,831 cows on 2,970 herds) classified by the three geographical areas (Po valley, Sub-Alps and Alps).

**Fig 6.**
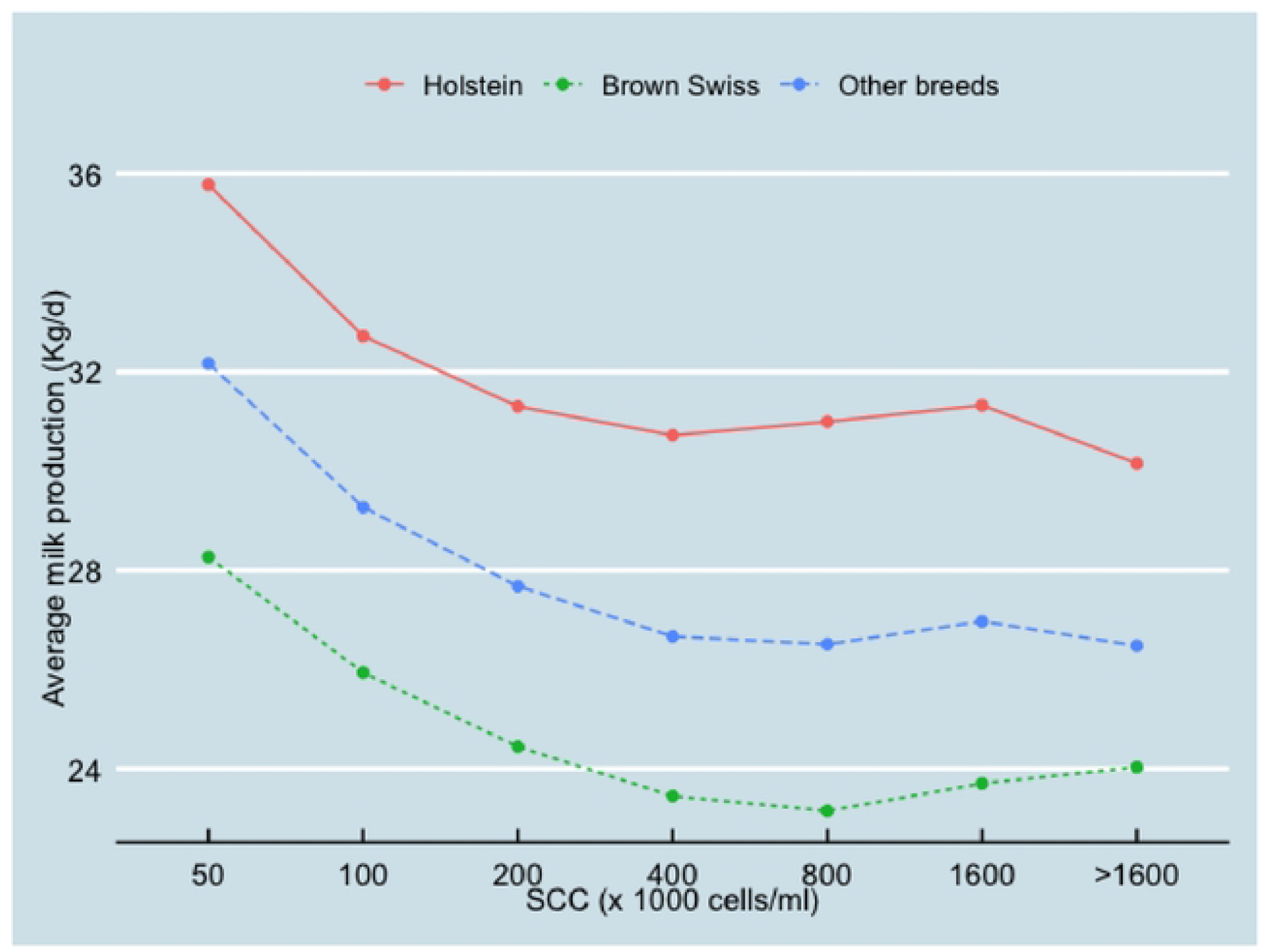
Relationship between SCC and average milk production(10,445,464 test-day records from 826,831 cows on 2,970 herds) classified by different breeds (Holstein, Brown Swiss and other breeds)

### Statistical analysis

The fixed effects estimate of the final linear mixed effect model on the association between SCC and milk production are presented in Table 2. The overall mean of milk production was 34.75 (95% confidence interval (CI): 34.59, 34.91) kg/cow/day at the reference level (in the Po valley area, Holstein as the breed, the first month of lactation, the first parity, autumn and 2016). For every unity of Log_2_SCC increase, the milk production decreased 0.83 (95% CI: −0.832, −0.828) kg/cow/day. In the Alps area, the milk production was the lowest. Holstein cows had the highest milk production among breeds, while Brown Swiss cows had the lowest milk production. Cows produced more milk in the second or higher parities than in their first parity. Autumn was the lowest milk production season. In each year, considered in this study, milk production increased gradually.

**Table 2.**
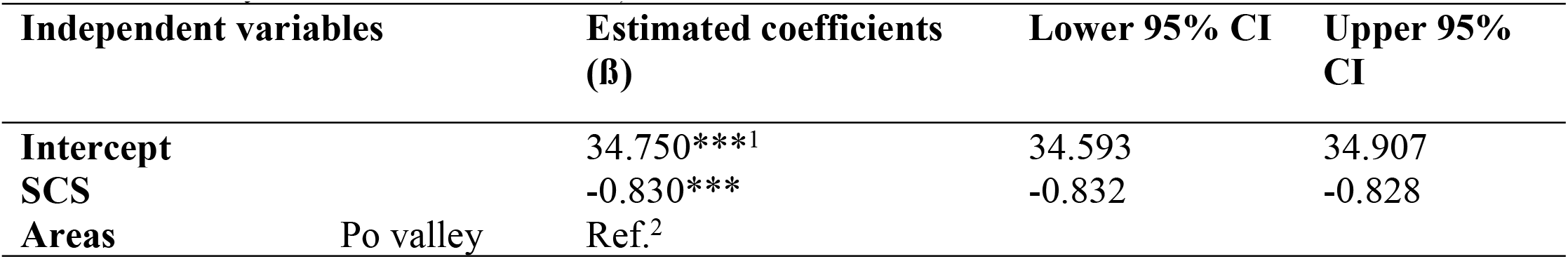
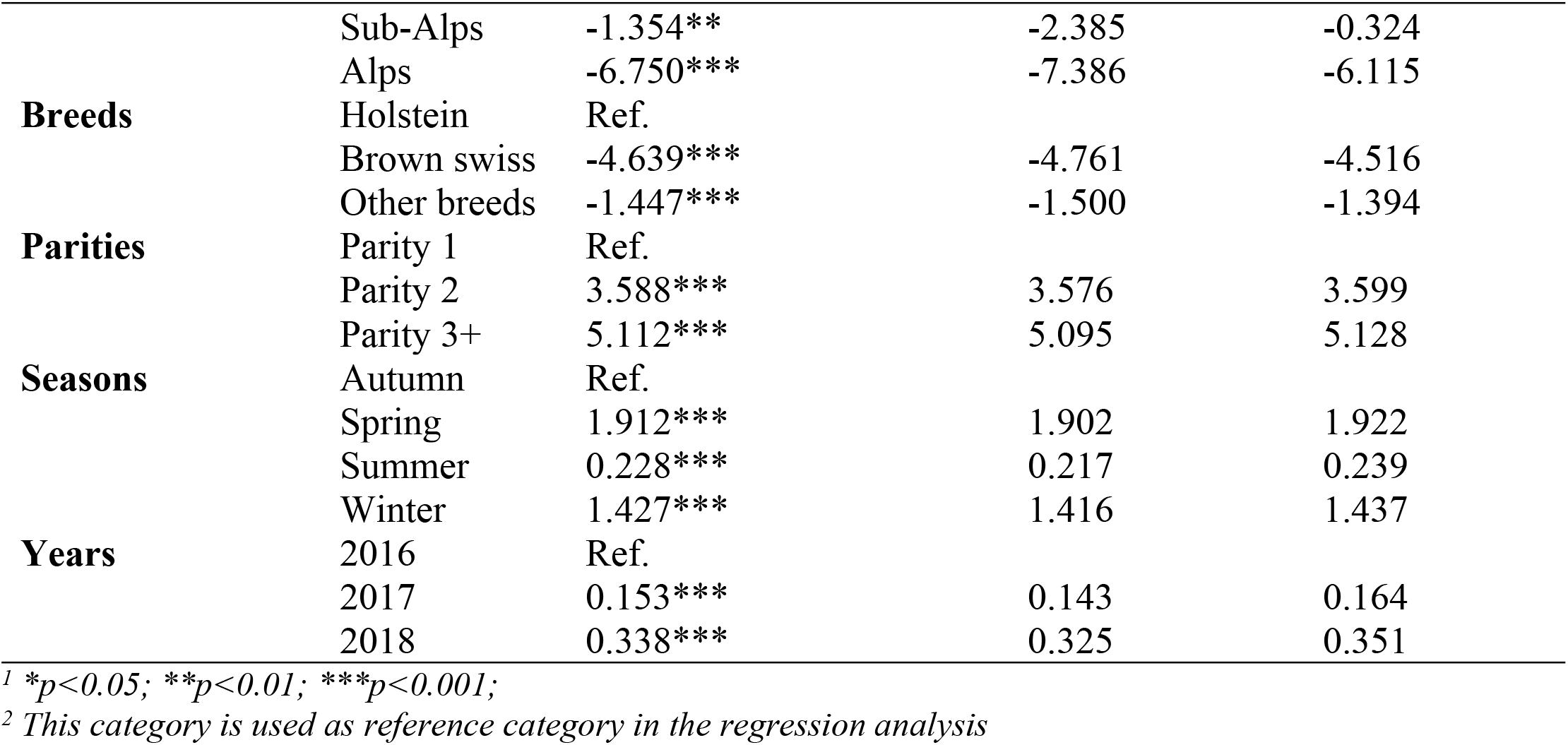
Coefficient estimates (β) of the linear mixed model for milk production (in kg/day). (The estimates for days in milk are not shown)

### The fit of the Model

The VIF of the fixed independent variables were all at the low level (lower than 1.25), indicated that the correlation between fixed independent variables was low. The marginal R-squared and the conditional R-squared were 0.31 and 0.67, respectively. The residual variability was explained by the random intercept in the model with an ICC of 0.52. No evidence indicates the presence of influential observations. From the visual inspection of the studentized residuals normal Q-Q plot, the normality of the residuals from the model was acceptable.

When the milk production was greater than 35 kg/day, there was an increasing non-linear relationship between predictor variables and the outcome variable. However, when the milk production was smaller 35 kg/day, the model was able to capture the linear relationship (see fig 7). In Fig 8, the square root of standard residuals and fitted values are plotted to check the assumption of homoscedasticity. There is an increasing level of heteroskedasticity when the fitted value is greater than 40 kg/day of milk.

**Fig 7.**
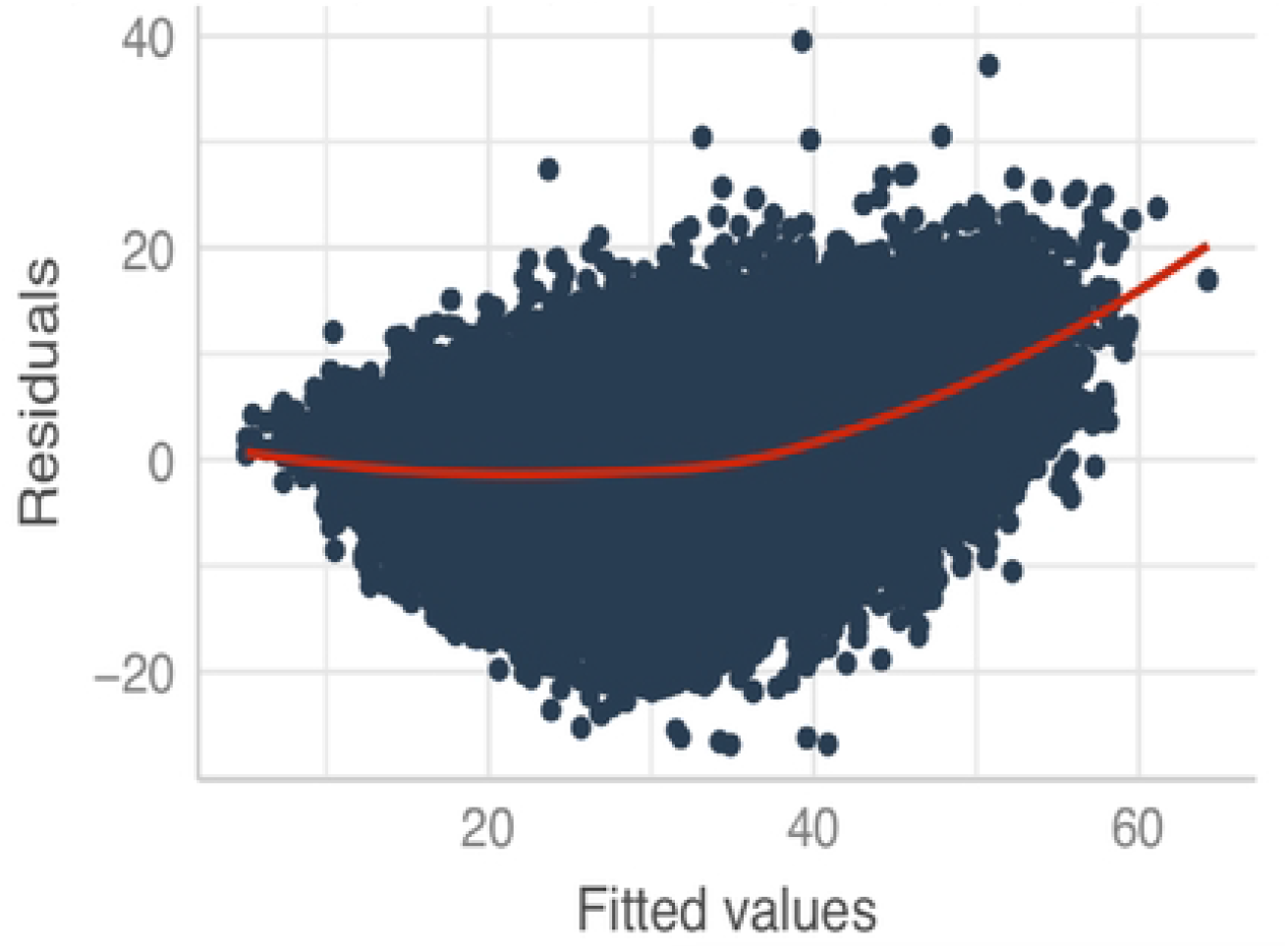
Standard residuals vs fitted values from the linear mixed model to estimate the association between SCC and milk production (10,445,464 test-day records).

**Fig 8.**
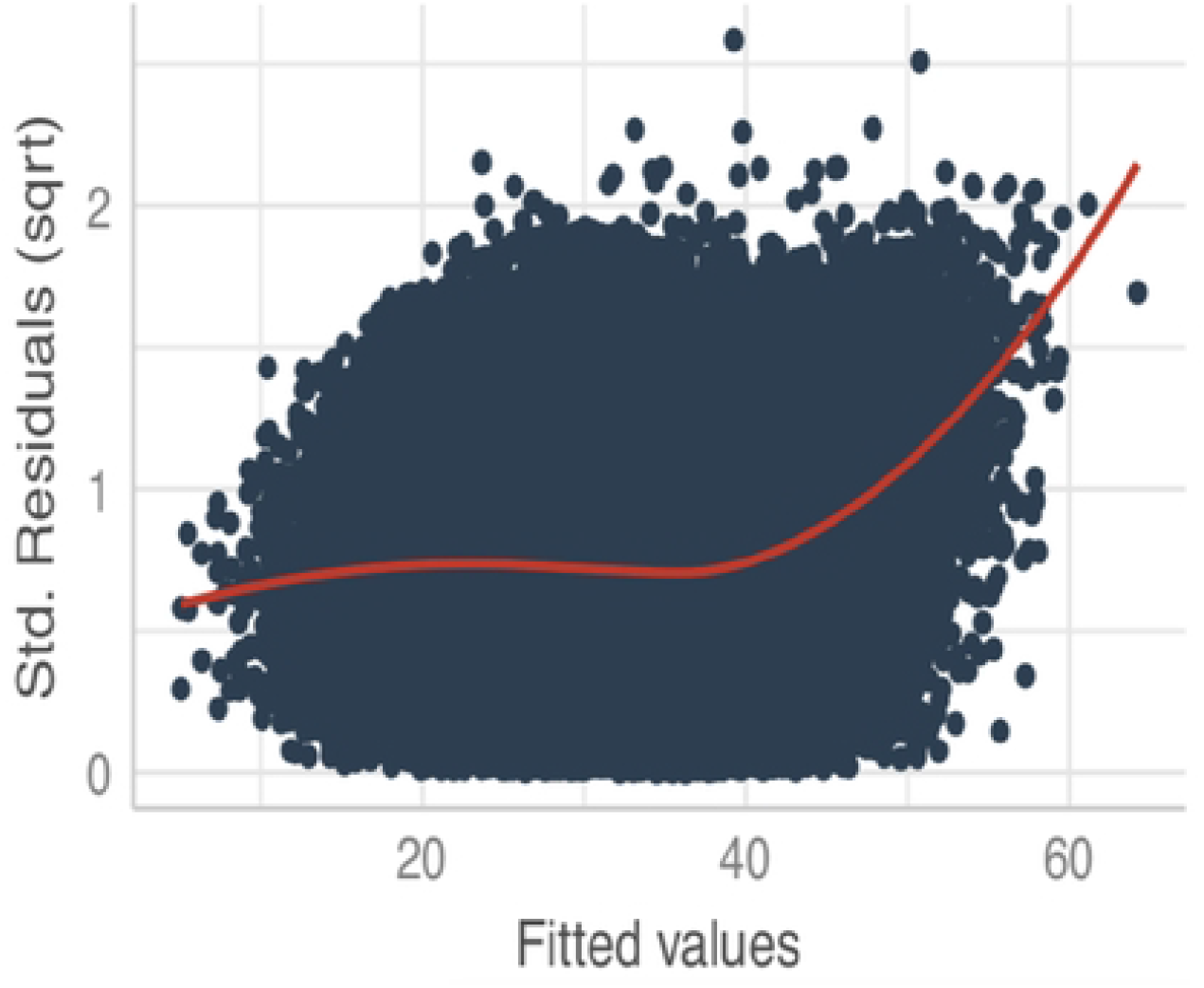
Scale-Location plot from the linear mixed model to estimate the association between SCC and milk production (10,445,464 test-day records).

## Discussion

In this study, the association between SCC and milk production was quantified for Lombardy dairy farms from 2016 to 2018. Results indicated that when the SCC increased every two-fold, the milk production lost 0.830 kg/cow/day in the whole Lombardy. The current study also revealed clear differences among the areas in Lombardy. In the Alps, the average herd size was the smallest (86 cows) with the highest average SCC (319,000 cells/mL) and the lowest average milk production (26.3 kg/d). In the Po valley, the average herd size was the largest (233 cows) with an average SCC of 308,000 cells/mL and the highest milk production (32.9 kg/d). The average herd size in the Sub-Alps was 109 cows, with an average SCC of 274,000 cells/mL and an averge milk production of 31.5 kg/d. The results of the final model confirmed these differences between regions. Compared with the Po valley (as the reference in the model), the coefficient estimates (β) of Alps and Sub-Alps were −1.354 kg/d and −6.750 kg/d, respectively. These differences may due to the different herd characteristics between Po valley, Sub-Alps and Alps dairy herds [22]. Despite the differences in the amount of milk production and the SCC level among areas and breeds, the presence of a strong association between these two parameters was found. Hence, Figures 5 and 6 showed that when SCC went from 50,000 cells/ml to 400,000 cells/ml milk production decreased with about 5 kg/d, independently from areas or breeds.

When the SCC increased every two-fold, the estimated decrease in milk production was higher in Lombardy than in other European countries. There, the decrease in milk production ranged from 0.2 kg/d to 0.5 kg/d with different mastitis conditions in the Netherlands, Sweden and other countries[13,17,23]. Moreover, in the current study, crude SCC was used to fit into the models, was not adjusted with the dilution effect [24], so the adjusted SCC could be higher than the observed values.

Also other factors might have influenced the results of the current study. Firstly, the average milk production and SCC in Lombardy were relatively high compared to other studies from different countries. For instance, the average milk production ranged from 23.2kg/d to 28.3 kg/d, and the average SCC ranged from 65,000 cells/mL to 105,000 cells/mL in the Netherlands[13,25]. The high milk production in Lombardy was due to several reasons. Lombardy dairy herds were genetically specialized with high milk production cows [26]. The farmers applied complex diets [27] by using the total mixed ration method perennially, without pasturing. Moreover, the farmers were highly motivated to achieve higher production because of the thin difference between feeding cost and revenues, thus a higher efficiency is needed to have a profit [28].

Secondly, Lombardy herds have different breeds, while other studies mainly analysed herds with only Holstein Friesian cows [18,29]. Cows in different breed groups had heterogeneous milk production and SCC levels. Holstein cows had the highest milk production and the lowest SCC among all breed groups, while Brown Swiss cows had the lowest milk production and the highest SCC among all breed groups (Figure 6). It’s worth noting that in different areas, the components of the categorical factor “other breeds group” were different. In the Alps area, a substantial number of the local breeds and Simmental were in this group, while in other Lombardy areas, the mixed breed with Holstein was the main component in the “other breeds” group. Since milk production capability of the local breeds and Simmental was smaller than the mixed breed with Holstein cows, in the Alps area, the cows in “other breeds group” had the lowest milk production and the highest SCC compared with Holstein cows and Brown Swiss cows.

Finally, autumn was the lowest milk production season in Lombardy (including the Alps area), which was due to the ambient temperature and the calving pattern. In Lombardy, all the herds applied similar calving pattern with cows starting to be dried off in the autumn which caused the lowest milk production. This finding was different from other studies [30,31] where summer or spring was the lowest production season.

The diagnosis plots of the model indicated that heteroscedasticity was present when the fitted values were higher than 40 kg/d. This means higher fitted values had larger residuals, and the model did not fit well with larger predicted means. Klein et al. (2016) revealed that the presence of heteroscedasticity could be due to missing some confounders or interaction terms in the model. However, in the current study, these potential confounders that may influence the milk production, for example, the feeds and farming technics, were controlled by the random herd effect in the models. Moreover, the different interaction terms, different random slopes and single parity data were checked by inserting them into the model, however, the heteroscedasticity was not solved. For simplifying the model and the purpose of this study these interaction terms and random slopes were not included in the final model.

Two potential consequences could be present due to heteroscedasticity: the least-squares estimator is still a linear and unbiased estimator, but with wider variance; and the standard errors computed for the least-squares estimators are incorrect[32]. This can affect confidence intervals and hypothesis testing that use those standard errors, which could mislead conclusions. However, due to the massive data size in the current study, the confidence intervals were all relevantly very small, this misclassification error should not happen. Furthermore, Schielzeth et al. [33]had proved the robustness of linear mixed-effects models. The results showed that the fixed effect estimates, in particular, were relatively unbiased when heteroscedasticity data were fitted in linear mixed-effects models. Thus, the association between SCC and milk production in Lombardia and the Alps area should be unbiased by using the mixed-effects model.

Some biases might occur in this study. The database was provided by the dairy farmer association in Lombardy. This might lead to selection bias since there is lacking randomness in the samples to represent all the dairy cows in Lombardy or Northern Italy. Moreover, this study excluded the small herds, which may lead to additional selection bias. This selection bias might have caused an underestimation of the loss of milk production since these small herds had higher SCC and lower milk production.

Furthermore, the assessment of this association in the Lombardy herds, suggested that applying values from studies performed in other countries may be misleading, since the drop in yield as SCC increases was higher. These information should be considered when the economic impact of subclinical mastitis (defined by SCC) is estimated to identify priorities in the application of herd management programs.

## Conclusion

This study focused on quantifying the association between SCC and milk production, by analyzing the TD testing records data of 2,970 Lombardy dairy herds from January 2016 to December 2018. The results confirmed that the SCC had a negative association with milk production. The analysis was carried out by using a mixed-effects model with seven fixed effects (geographical area, breed, days in milk, parity, season and year) and nested random effects for each cow and herd. The results indicated that the SCC had a negative association with milk production. When the SCC increased every two-fold, the milk production lost 0.830 (95% CI: −0.832, −0.828) kg/cow/day in the whole Lombardy. The pattern of the relationship between SCC and milk production showed when SCC went from 50,000 cells/ml to 400,000 cells/ml milk production decreased with about 5 kg/d, independently from areas or breeds. The results suggest that an improvement in herd udder health will result in a significant increase in milk yield and, therefore, of herd efficiency.

## References

1. T. Halasa, K. Huijps OØ& HH. Economic effects of bovine mastitis and mastitis management: A review. Vet Q. 2007;29: 18–31. doi:10.1080/01652176.2007.9695224

2. Hogeveen H, Huijps K, Lam TJGM. Economic aspects of mastitis: New developments. N Z Vet J. 2011;59: 16–23. doi:10.1080/00480169.2011.547165

3. Ruegg PL, Petersson-Wolfe CS. Mastitis in Dairy Cows. Vet Clin North Am - Food Anim Pract. 2018;34: ix–x. doi:10.1016/j.cvfa.2018.08.001

4. Hogeveen H, Steeneveld W, Wolf CA. Production Diseases Reduce the Efficiency of Dairy Production: A Review of the Results, Methods, and Approaches Regarding the Economics of Mastitis. Annu Rev Resour Econ. 2019;11: 289–312. doi:10.1146/annurev-resource-100518-093954

5. Hadrich JC, Wolf CA, Lombard J, Dolak TM. Estimating milk yield and value losses from increased somatic cell count on US dairy farms. J Dairy Sci. 2018;101: 3588–3596. doi:10.3168/jds.2017-13840

6. Huijps K, Lam TJGM, Hogeveen H. Costs of mastitis: Facts and perception. J Dairy Res. 2008;75: 113–120. doi:10.1017/S0022029907002932

7. van Soest FJS, Santman-Berends IMGA, Lam TJGM, Hogeveen H. Failure and preventive costs of mastitis on Dutch dairy farms. J Dairy Sci. 2016;99: 8365–8374. doi:10.3168/jds.2015-10561

8. Zecconi A ZG. Il controllo delle mastiti per un allevamento sostenibile. Bologna: Filiera AQ - Università degli Studi di Milano; 2013.

9. Zecconi A, dell’Orco F, Rizzi N, Vairani D, Cipolla M, Pozzi P, et al. Cross-sectional study on the prevalence of contagious pathogens in bulk tank milk and their effects on somatic cell counts and milk yield. Ital J Anim Sci. 2020;19: 66–74. doi:10.1080/1828051X.2019.1693282

10. Alhussien MN, Dang AK. Milk somatic cells, factors influencing their release, future prospects, and practical utility in dairy animals: An overview. Veterinary World. 2018. doi:10.14202/vetworld.2018.562-577

11. Farre, M.; Zecconi, A.; Kelton D. Guidelines for defining quarter and udder health status and cured clinical and subclinical mastitis cases. IDF Bulletin 515, Bruxelles; 2022.

12. Zecconi A, Vairani D, Cipolla M, Rizzi N, Zanini L. Assessment of subclinical mastitis diagnostic accuracy by differential cell count in individual cow milk. Ital J Anim Sci. 2019;18: 460–465. doi:10.1080/1828051X.2018.1533391

13. Halasa T, Nielen M, De Roos APW, Van Hoorne R, De Jong G, Lam TJGM, et al. Production loss due to new subclinical mastitis in Dutch dairy cows estimated with a test-day model. J Dairy Sci. 2009;92: 599–606. doi:10.3168/jds.2008-1564

14. Hand KJ, Godkin A, Kelton DF. Milk production and somatic cell counts: A cow-level analysis. J Dairy Sci. 2012;95: 1358–1362. doi:10.3168/jds.2011-4927

15. Juliano L. Gonçalves, Roger I. Cue, Bruno G. Botaro, José A. Horst, Altair A. Valloto, Santos and M V. Milk losses associated with somatic cell counts by parity and stage of lactation. J Dairy Sci. 2018;101: 4357–4366. doi:10.3168/jds.2017-13286

16. Chen H, Weersink A, Kelton D, von Massow M. Estimating milk loss based on somatic cell count at the cow and herd level. J Dairy Sci. 2021. doi:10.3168/jds.2020-18517

17. Hagnestam-Nielsen C, Emanuelson U, Berglund B, Strandberg E. Relationship between somatic cell count and milk yield in different stages of lactation. J Dairy Sci. 2009;92: 3124– 3133. doi:10.3168/jds.2008-1719

18. Gonçalves JL, Cue RI, Botaro BG, Horst JA, Valloto AA, Santos M V. Milk losses associated with somatic cell counts by parity and stage of lactation. J Dairy Sci. 2018;101: 4357–4366. doi:10.3168/jds.2017-13286

19. Kirk JH. Programmable Calculator Program for Linear Somatic Cell Scores to Estimate Mastitis Yield Losses. J Dairy Sci. 1984;67: 441–443. doi:10.3168/jds.S0022-0302(84)81322-3

20. Nieuwenhuis R, te Grotenhuis M, Pelzer B. Influence.ME: Tools for detecting influential data in mixed effects models. R J. 2012;4: 38–47. doi:10.32614/rj-2012-011

21. Bates D, Mächler M, Bolker BM, Walker SC. Fitting linear mixed-effects models using lme4. J Stat Softw. 2015;67. doi:10.18637/jss.v067.i01

22. Penati CA, Tamburini A, Bava L, Zucali M, Sandrucci A. Environmental impact of cow milk production in the central Italian Alps using Life Cycle Assessment. Ital J Anim Sci. 2013;12: 584–592. doi:10.4081/ijas.2013.e96

23. Henri SEEGERS, Christine FOURICHON FB. Production effects related to mastitis and mastitis economics in dairy cattle herds. Vet Res. 2003;34: 475–491. doi:10.1051/vetres

24. Green LE, Schukken YH, Green MJ. On distinguishing cause and consequence: Do high somatic cell counts lead to lower milk yield or does high milk yield lead to lower somatic cell count? Prev Vet Med. 2006;76: 74–89. doi:10.1016/j.prevetmed.2006.04.012

25. Yue X, Steeneveld W, van der Voort M, van Schaik G, Vernooij JCM, van Duijn L, et al. The effect of bovine viral diarrhea virus introduction on milk production of Dutch dairy herds. J Dairy Sci. 2021;104: 2074–2086. doi:10.3168/jds.2020-18866

26. Macciotta NPP, Mele M, Conte G, Serra A, Cassandro M, Dal Zotto R, et al. Association between a polymorphism at the stearoyl CoA desaturase locus and milk production traits in Italian holsteins. J Dairy Sci. 2008;91: 3184–3189. doi:10.3168/jds.2007-0947

27. Abeni F, Calamari L, Stefanini L, Pirlo G. Effect of average daily gain on body size, metabolism, and milk production of Italian Holstein heifers raised on two different planes of nutrition and calving at two different ages. Livest Sci. 2012;149: 7–17. doi:10.1016/j.livsci.2012.06.003

28. Bellingeri A, Cabrera V, Gallo A, Liang D, Masoero F. A survey of dairy cattle management, crop planning, and forages cost of production in Northern Italy. Italian Journal of Animal Science. 2019. pp. 786–798. doi:10.1080/1828051X.2019.1580153

29. Wilson DJ, González RN, Hertl J, Schulte HF, Bennett GJ, Schukken YH, et al. Effect of clinical mastitis on the lactation curve: A mixed model estimation using daily milk weights. J Dairy Sci. 2004;87: 2073–2084. doi:10.3168/jds.S0022-0302(04)70025-9

30. Liang D, Wood CL, McQuerry KJ, Ray DL, Clark JD, Bewley JM. Influence of breed, milk production, season, and ambient temperature on dairy cow reticulorumen temperature. J Dairy Sci. 2013;96: 5072–5081. doi:10.3168/jds.2012-6537

31. Ray DE, Halbach TJ, Armstrong D V. Season and Lactation Number Effects on Milk Production and Reproduction of Dairy Cattle in Arizona. J Dairy Sci. 1992;75: 2976–2983. doi:10.3168/jds.S0022-0302(92)78061-8

32. Long JS, Ervin LH. Using Heteroscedasticity Consistent Standard Errors in the Linear Regression Model. Am Stat. 2000;54: 217–224. doi:10.1080/00031305.2000.10474549

33. Schielzeth H, Dingemanse NJ, Nakagawa S, Westneat DF, Allegue H, Teplitsky C, et al. Robustness of linear mixed-effects models to violations of distributional assumptions. Methods Ecol Evol. 2020;11: 1141–1152. doi:10.1111/2041-210X.13434

